# Illuminance affects epidemiological parameters of banana Yellow Sigatoka in Brazil

**DOI:** 10.1101/828848

**Authors:** Djalma M. Santana-Filho, Milene C. da Silva, Jorge T. de Souza, Zilton J. M. Cordeiro, Hermínio S. Rocha, Francisco F. Laranjeira

## Abstract

The Sigatoka leaf spots are among the most important banana diseases. Although less damaging than black sigatoka, yellow sigatoka (*Pseudocercospora musae*) still prevails in some regions. This study aimed at testing the hypothesis of light interference in monocyclic parameters of yellow sigatoka epidemics. Grande Naine plantlets kept under contrasting shading conditions had their leaves 1 and 2 inoculated. Evaluations were performed for 60 days. For each inoculated leaf, the time until symptom onset (incubation), presence of infectious lesions (latency), and disease severity (extensive leaf necrosis) according to Stover’s scale modify per Gauhl (1994), called here only Stover’s scale, were registered. Logistic regression was used to assess the relative occurrence risk and survival analysis was used to check the effects of variables on relevant epidemiological parameters. The risks of sporulation and of reaching high severities were lower for plants kept under shading regardless of the acclimation conditions and no effect of leaf age was detected. The logistic regression showed symptoms appearing in both conditions (p=0,85), but have significance difference in occurrence of latent lesions (p=0,013) and necrosis (p<0,0001). The necrosis risk in non-shaded environment arrived 66%. The survival analysis showed significance difference in the time to appear the symptom evaluated in all tested variables (p<0,0001) in function of the cropping system. Lower illuminance negatively affected the incubation, latency and infectious periods, and severity. A shaded system could be tested to produce organic bananas in areas of high risk of occurrence of Yellow sigatoka disease.

**Significance and Impact of the Study:** Yellow Sigatoka (*Pseudocercospora musae*) is a banana disease that can cause severe damage if left uncontrolled. Its control is based mostly on fungicides.Our results show that shading downregulates the epidemiological parameters of that disease such as incubation, latent and infectious periods, and symptom’s severity. These results can be the basis for testing alternative cropping systems and producing organic bananas.

## INTRODUCTION

Sigatoka leaf spot (popularly known as Yellow Sigatoka) is a *Musa* sp. fungal disease caused by *Pseudocercospora musae* (syn. *Mycosphaerella musicola*) (Carlier et al., 1994). This disease is found in most banana growing regions of the world (Mourichon, 2002; Crous et al., 2003) and although is not more severe than black Sigatoka (black leaf streak) caused by *P. fijiensis* (syn. *M. fijiensis*) (Mourichon, 2002; Fullerton, 1994; Jones, 2000 apud Marín et al., 2003; Churchill, 2011), it can still cause considerable losses at higher altitudes and cooler temperatures, and is also typically a greater problem during rainy seasons in subtropical banana growing regions (Mouliom-Pefoura et al. 1996). The disease reduces the leaf’s photosynthetic capacity, which affects bunch size. It also shortens the fruit’s green life and the time between harvest.

Despite both pathogens occurring in Brazil, *P. musae* is more disseminated and responsible to decrease the banana yield in the field (Gomes et al., 2013: Silveira et al., 2016). The pathogen infect through stomata, mainly on the three younger leaves, although infection may occur on the fourth leaf in case of severe outbreaks (Agrios, 2005). The first symptoms appear three to four weeks after spores land on susceptible leaf surfaces (Hoss et al., 2000). These diseases have an endemic nature in tropical countries and epidemics may occur depending on local climatic conditions and the adopted crop management (Fourè, 1994). The main environmental factors influencing infection, inoculum production and dissemination are temperature, rain and dew (Cordeiro et al., 2004). These factors can also affect the incubation period (Cordeiro, 1997), the latent period (Marin et al., 2003) and the development of lesions (Fourè, 1994).

Banana cultivation in a semi-shaded scheme, as agroforestry systems or under high density, is thought to be an alternative cropping system to manage Yellow Sigatoka. Studies on black Sigatoka show that for plants grown under nylon shading meshes (30% light) there is a significant decrease in the disease’s incidence and a slower disease development than at 70% of light (Norgrove, 1998; Favreto et al., 2007; Dold et al., 2008; Beltrán-García et al., 2014). In low-density plantations (1600-2500 plants/ha), removal of old infected leaves leads to a 16% reduction in disease incidence and 10% in severity, but at high densities (3333 plants/ha), where there is more shade, incidence was not affected and severity was reduced by 18% (Emebiri & Obiefuna, 1992). This shows that the light level could be interfering in the fungal life cycle.

The lower black Sigatoka severity registered under shade conditions was hypothesized to correlate with a diminished activation of the pathogen’s toxin cercosporin and lower amounts of reactive oxygen species generated by melanin in the shade (Daub et al., 2005; Beltrán-García et al., 2014). Furthermore, the light also has been shown to influence the *P. fijiensis* sporulation (Sepúlveda et al., 2009).

Shade is also supposed to limit damages caused by *P. musae* (Vivan, 2002), but no detailed studies have been carried out for yellow Sigatoka. Soon, this is an excellent opportunity to study the relation between the illuminance, amount of light per unit area, and the Yellow Sigatoka life cycle.

The objective of this study was to assess light interference with epidemiological parameters of yellow Sigatoka. A hypothesis of light-dependent Sigatoka was tested by characterizing the risks of infection, sporulation and leaf necrosis in inoculated plants. The hypothesis of light-enhanced Sigatoka was tested by quantifying the length of incubation, latency, and infectious periods. The light-dependent hypotheses admit that the variables occurs in the illuminated environmental and not in the shade. On the other hand, the light-enhanced hypotheses admit that the variable occurs in two situations, illuminated or shaded environmental, and that the light makes increase and shade decrease in the time of development of disease.

## MATERIALS AND METHODS

Banana plantlets (before *Musa paradisiaca* L. var. Grand Naine, now *Musa acuminate* L. var. Grande Naine) were produced in vitro (Campo, Cruz das Almas, Bahia). Then, half of plantlets were acclimated at full sun and the other half under 50% light.

After 130 days of cultivation the plantlets were taken to shade house to inoculation and start the experiment. The shade house was 2m high and covered with a dark nylon screen (Sombrite 75%). The roof was fixed, but the sides were built with rolling screens. The side screens were rolled up every night and in periods during the day when sunlight could not directly reach the plants. Hence, minimizing differences in temperature, relative humidity and leaf wetness between shaded and non-shaded areas. Before inoculation, stomata density was determined right before fungal inoculations by pressing a glass slide covered with a thin layer of Loctite adhesive on the leaves surface for 2”. The adhesive layer was examined under a microscope at 10X and the number of stomata per area determined.

The inoculation was done with the isolate CNPMF 06/10 of *P.* musae, obtained from the municipality of Muritiba, Bahia and deposited in the Embrapa Cassava & Fruits culture collection, was used in all inoculations. Spores were produced according to Cordeiro et al (2011). Spore suspensions were adjusted to 4 × 10^4^ conidia/mL and used to spray leaves 1 and 2 until run-off; non-inoculated plants were sprayed with sterile water. The numbering of the banana leaves starts from the youngest leaf, with the number zero, which is still curled, followed by the older leaves 1, 2, and so on, which are already totally Unrolled. Inoculations took place at dusk to prevent illuminance effects in the pre-penetration processes. All plants were covered with plastic bags after inoculation and kept under shading overnight. The non-inoculated leaves was used to check the natural infections from environmental.

Fourteen hours after the inoculation the plastic bags were removed. The plantlets were arranged in completely randomized blocks, with four groups: T1) plantlets kept under full sun and acclimated in the sun; T2) plantlets kept under sun and acclimated in the shade; T3) plantlets kept under shade and acclimated in the sun, and T4) plantlets kept in the shade and acclimated in the shade. Due to plant losses during acclimation, the experimental groups were different for each treatment. T1 and T3 had 60 inoculated and 40 non-inoculated plantlets; T2 had 60 inoculated and 19 non-inoculated plantlets, and T4 had 40 inoculated and 19 non-inoculated plantlets. Groups T3 and T4 we kept in the shade house whereas the other two were kept in an adjoining area the same size as the shaded one.

Every eight days disease severity was evaluated with the aid of a scale described by Stover modify per Gauhl (1994), where (0) represents no symptoms; (1) less than 1% of the leaf tissue lesioned or a maximum of 10 lesions; (2) 1 to 5% of lesioned leaf tissue; (3) 6 to 15% of lesioned area; (4) 16 to 33%; (5) 34 to 50%; (6) 51 to 100%; (-) dead necrotic leaf hanging on the pseudostem. Leaves number 1 and 2 were evaluated daily for the following variables: occurrence of the first lesion (incubation period); occurrence of the first sporulating lesion (latent period) observing gray lesions, number of days to reach disease index 4 according to the scale (severity), number of days to reach the dash index (necrosis). The infectious period was defined as the time between latency and necrosis and illuminance as the amount of light casting in a determined area. Three explanatory variables were tested: (1) shade conditions during acclimation; (2) shade conditions after inoculation; (3) age of the inoculated leaf (number 1 or 2).

The illuminance (lux, luminous power incident on a surface) was measured daily at noon in at least four positions of each contrasting areas using a photometer. Temperature was also registered daily in shaded and non-shaded areas.

Due to the longitudinal nature of the experiment, different groups sizes, and occurrence of censored data, we used survival analysis (Statistical release program 7.1) and logistic regression as statistical tools. The logistic regression (BioEstat 5.0) was applied to model the risk imposed by the explanatory variables to the occurrence of an event. In this case, both explanatory and dependent variables are binary [e.g. shade (0) or no-shade (1) versus no symptoms (0) or symptoms (1)]. The events of interest were the presence of symptoms, the presence of sporulating lesions, occurrence of high severities (level 4 in the Stover’s scale) and occurrence of extensive leaf necrosis (level 6). The coefficients of each regression were calculated using maximum likelihood. A given model was accepted only when the regression and the coefficients were statistically significant (p<0.05). The odds ratio (probability of occurrence: probability of non-occurrence) was calculated for each significant explanatory variable.

Survival analysis was used to model the time to the events of interest. Here, we considered the incubation period, the latency period, and the time to disease development, to high severities and to necrosis, as defined above. The proportional hazards model was fitted to the data, and its adequacy was tested by χ^2^ test. A model was accepted only when the coefficients of the explanatory variables were significant (p<0.05). Hazards ratios werecalculated for each of them. Kaplan-Meyer curves (Statistical release 7.1) were constructed for each combination of significant explanatory variables, and used to estimate the median expectancy of each epidemiological period of interest.

## RESULTS

### Illuminance, Temperature and Stomatal density

The illuminance under shaded conditions was approximately 26% of that registered at the non-shaded area (Figure 1). The mean of stomatal density of plants acclimated under shading was 38.5±6.4 stomata per mm^2^ whereas it was 50.8± 8.5 stomata per mm^2^ for those acclimated under the sun. The temperature under the shading apparatus was 30.1±2.5 and 27.6±1.1 at the non-shaded area. The difference in the median values to both stomata and temperature were significant (p <0.001) according the Mann-Whitney statistical test. In the shaded area we observed lower number of stomata/mm^2^ and higher temperature averages, while in the sunny area we had more stomata/mm^2^ and lower temperature averages.

**Figure 1.**
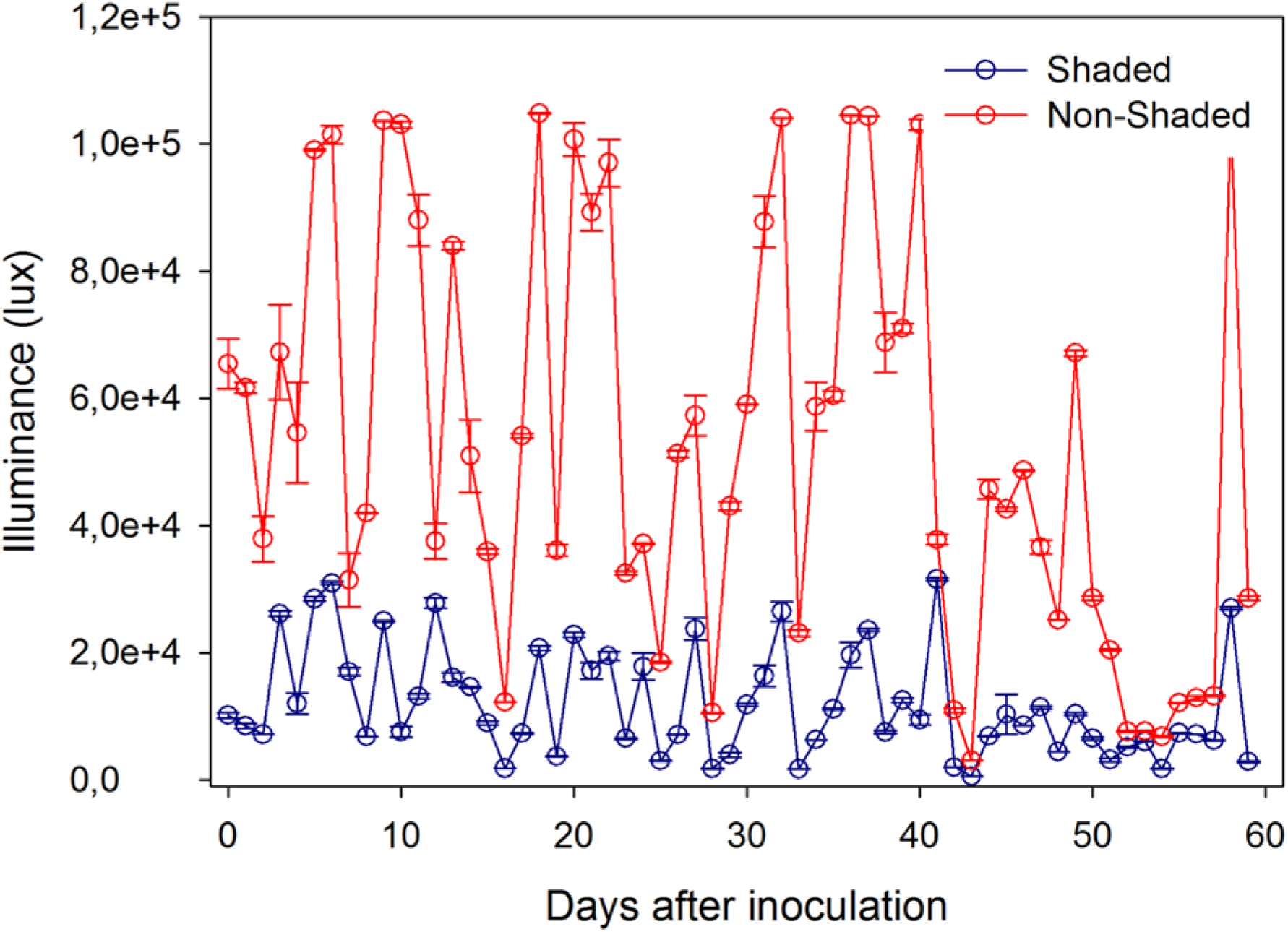
Illuminance values ± standard error along the experiment for the shaded and the non-shaded experimental areas.

### Experimental data

The inoculated leaves maintained under full sun had the time of incubation, latency, and to reach severity 4 and 6 less than those under shade (Table 1). All epidemiological parameters evaluated occurred earlier in plants grown in the sun than in plants grown in the shade (Table 2). The percentage of plants that completed their incubation period at the end of the experiment was equal, but these percentages were higher for plants completing the latency and infectious periods, and for plants that reached severity 4 and 6 (Table 2). These data were subjected to modelling with two different approaches: logistic regression and survival analysis.

**Table 1.**
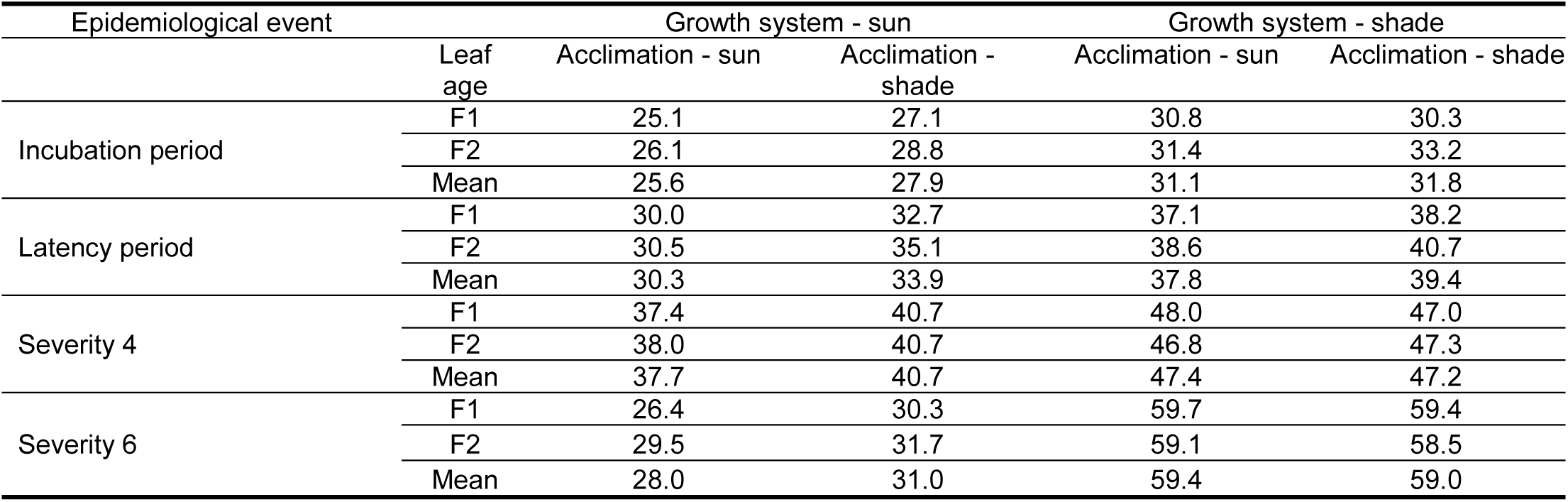
Average number of days for the occurrence of epidemiological events in plants acclimated and grown in the sun or in the shade.

| Epidemiological event | Leaf age | Growth system - sun |  | Growth system - shade |  |
| --- | --- | --- | --- | --- | --- |
|  |  | Acclimation - sun | Acclimation - shade | Acclimation - sun | Acclimation - shade |
| Incubation period | F1 | 25.1 | 27.1 | 30.8 | 30.3 |
|  | F2 | 26.1 | 28.8 | 31.4 | 33.2 |
|  | Mean | 25.6 | 27.9 | 31.1 | 31.8 |
| Latency period | F1 | 30.0 | 32.7 | 37.1 | 38.2 |
|  | F2 | 30.5 | 35.1 | 38.6 | 40.7 |
|  | Mean | 30.3 | 33.9 | 37.8 | 39.4 |
| Severity 4 | F1 | 37.4 | 40.7 | 48.0 | 47.0 |
|  | F2 | 38.0 | 40.7 | 46.8 | 47.3 |
|  | Mean | 37.7 | 40.7 | 47.4 | 47.2 |
| Severity 6 | F1 | 26.4 | 30.3 | 59.7 | 59.4 |
|  | F2 | 29.5 | 31.7 | 59.1 | 58.5 |
|  | Mean | 28.0 | 31.0 | 59.4 | 59.0 |

**Table 2.** Days for the onset of the epidemiological events and percentage of plants with symptoms at 60 days to sun and shade conditions.

| Event | Days for the onset of the epidemiological events |  | Percentage of plants with symptoms at 60 days |  |
| --- | --- | --- | --- | --- |
|  | Growth system |  | Growth system |  |
|  | Sun | Shade | Sun | Shade |
| Incubation period | 20 | 26 | 99 | 99 |
| Latency period | 23 | 33 | 99 | 90 |
| Severity 4 | 36 | 43 | 90 | 58 |
| Severity 6 | 43 | 52 | 79 | 5 |
| Infectious period | - | - | 89 | 6 |

### Light-dependent hypothesis

The logistic regression determines the relationship between the occurrence of one epidemiological event and the independent variables at the end of the experiment. It shows the chance of leaves showing an event under one or another system. The logistic regression was not significant for the incubation period when the variables acclimation, growth system and leaf number were considered (p=0.85). Likewise, latent period, time to severity 4 and time to severity 6 were not significant when the model considered the effect of the variables acclimation, growth system and leaf number together. However, a simpler model that took into account only the variable growth system was significant (Table 3). The latent period for plants grown in the sun have a risk 4X higher to produce sporulated lesions than plants grown in the shade (Table 3).

**Table 3.**
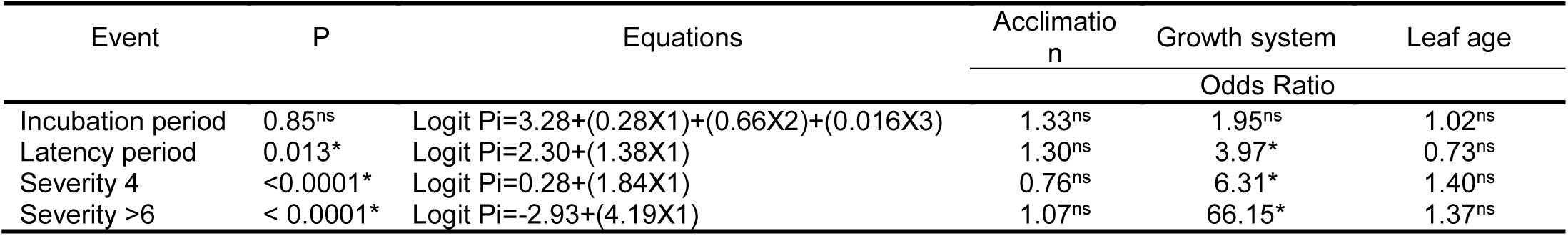
Logistic regression showing significance (P), and the significance and odds ratio of each explanatory variable for two events related to epidemiological parameter (occurrence of symptoms and sporulating lesions) and two severity related events. Leaf type and pre-inoculation conditions were non-significant whereas post-inoculation conditions was significant for all but the occurrence of symptoms. Leaves from plants kept non-shaded had more than 6x the chance of reaching Stover 4 than those on shaded plants.

The effect of growth system was significant for the occurrence of lesions on 16-33% of the leaf area (severity index 4 according to Gauhl’s scale) and necrosis (severity index 6) according to the multiple logistic regression (p<0.0001). The risk of occurring severity index 4 was 6X higher for non-shaded plants. Considering the occurrence of severity index 6, the risk was 66X higher for non-shaded plants (Table 1).

### Light-enhanced hypothesis

The survival analysis determines the effect of the explanatory variables at each day until the occurrence of the event of interest. Such analysis was used to quantify the delay in the epidemiological periods. The interference of illuminance on incubation and latency periods was significant for acclimation and growth system according to Cox’s model of proportional risks (Table 4). The probabilities of a shorter incubation period (1.3X) and a shorter latency period (1.4X) are higher for non-shaded acclimated plants. The risk of shorter incubation period (2.2X) and a shorter latency period (2.5X) are higher for plants grown in non-shaded area (Table 4).

**Table 4.**
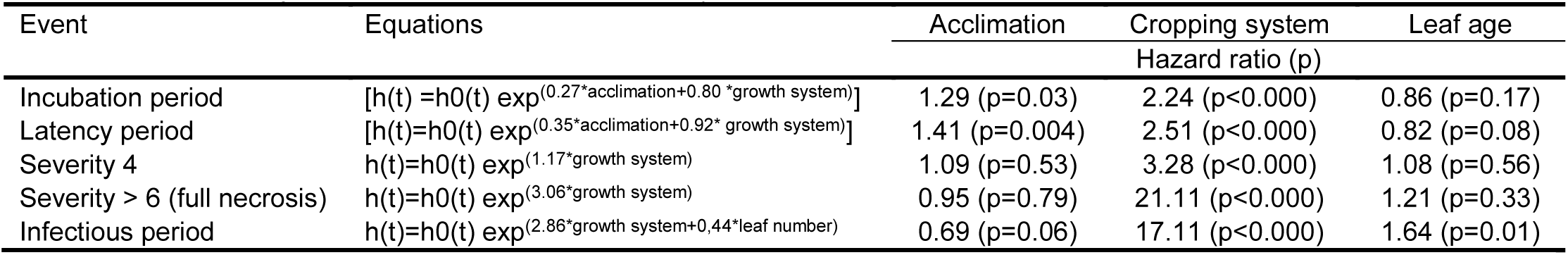
Results of Cox’s proportional hazards regression (survival analysis) showing the significance and the estimated hazard ratio of each explanatory variable for three epidemiological parameters (incubation, latency and infectious periods) and two severity events. Leaf type had no effect on the time until the events, whereas pre-inoculation conditions affected only the latency period. Post-inoculation conditions affected the times to all events (e.g. non-shaded leaves reach Stover 4 severity 3.28x faster than shaded ones).

The Kaplan-Meyer (KM) curves show the real data with the adjusted models and allow the medians to be calculated. Treatments combining sun-sun (acclimation-growth system), shade-sun and sun-shade, 90% of the plants showed symptoms before 42 days after the inoculation, whereas treatments combining shade-shade the first symptoms appeared after 51 days (Figure 2A). At the end of the experiment, 80% of the shade-shade (acclimation-growth system) plants and 90% of sun-shade and sun-sun plants showed sporulated lesions according to the KM curves (Figure 2B).

**Figure 2.**
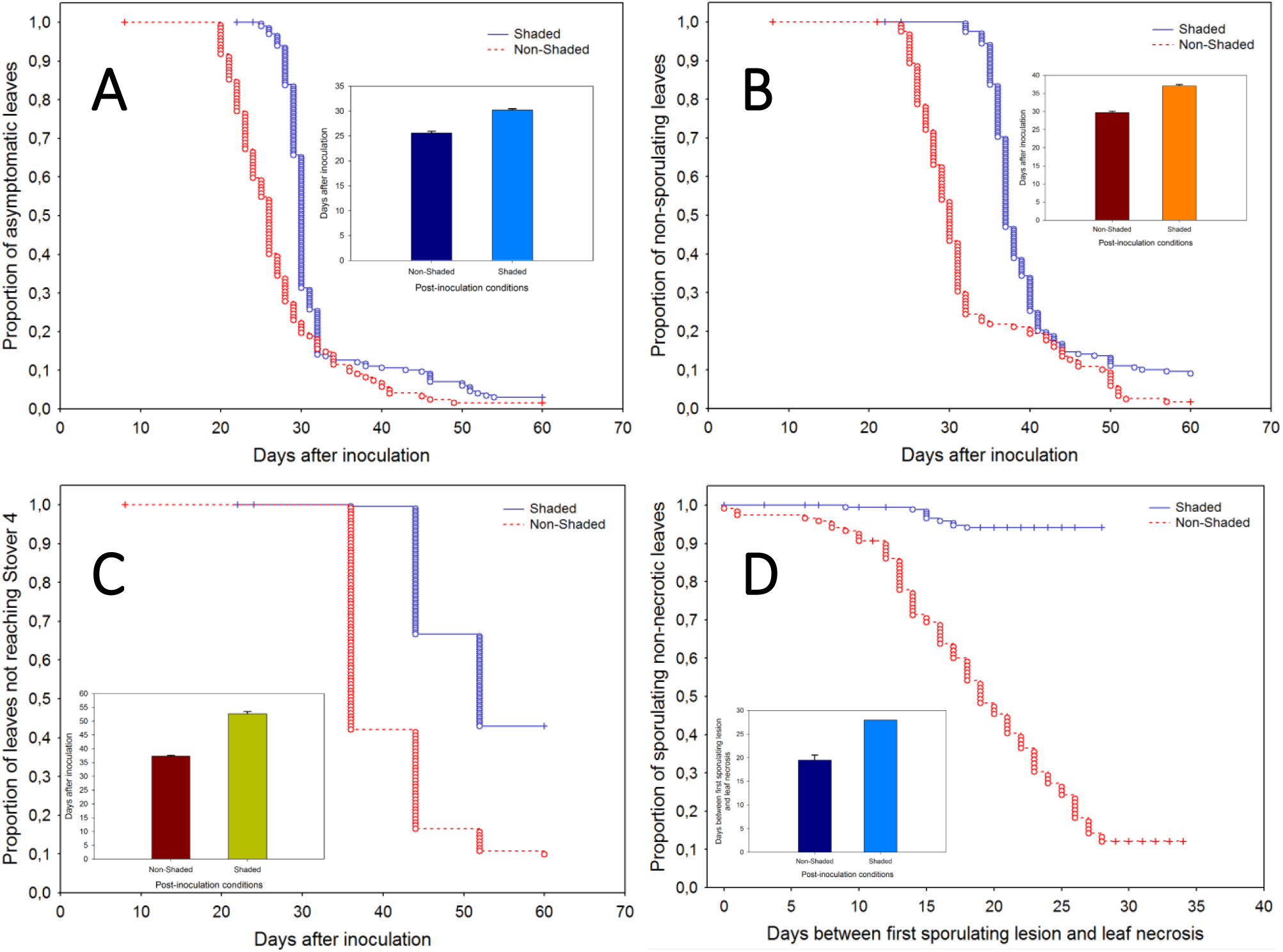
Kaplan-Meyer curves under shaded (blue) and non-shaded (red) post-inoculation conditions for incubation period (A), latency period (B), time to severity Stover 4 (C) and infectious period (D) of yellow Sigatoka. The insets show the median expected time ± standard error of each parameter under the two post-inoculation conditions. Circles represent complete observations and the crosses represent the censored observations. As less the half of the shaded leaves were necrotic at experiment completion, the related infectious period estimate is not precise. Note that the difference between shaded and non-shaded conditions is progressive (incubation: 4.6 days, latency: 7.4 days, Stover 4: 15.5 days). The results are consistent with the hypothesis of light-activated phytotoxins affecting the colonization process. On the other hand, the shorter latency on non-shaded leaves can influence the reproduction process.

There was an influence of the growth conditions on the time to reach severity 4 and severity 6 (necrosis). According to Cox’s model of proportional risks, non-shaded have a 3.3X higher chance of showing lesions on 16-33% of the leaf area than shaded grown plants. Similarly, the chance of showing more than 50% of lesioned leaf area is 21X higher for non-shaded grown plants (Table 4).

The KM curves showed that 36 days was required for the first non-shaded grown plants to present severity 4, whereas it took 44 days for shaded plants. The model for growth system (Figure 2D) shows that under shade 10% of the plants had the infectious period defined and 90% of the plants grown in the sun had lesions with the infectious period defined.

The infectious period was influenced by growth system (p<0.0001) and leaf age (p=0.015), but only marginally by acclimation (p=0.064). The model of proportional risks of Cox, showed a risk 17X higher of occurring the infectious period for non-shaded grown plants and 1.6X higher for leaves number 2 than for leaves number 1.

## DISCUSSION

The methodologies we used to study yellow Sigatoka disease were able to explain its behaviour in shaded and non-shaded systems. With the logistic regression analysis we calculate the risk of the pathogen present the incubation, latency or different levels of severity on shaded or non-shaded systems. The survival analysis was used to quantify the delay on the incubation, latency and different levels of severity. The first analysis was used to check if the variable occurs or not in function of illuminance, and the second one to compare and explain the behaviour of the variable, when it appear in shaded and non-shaded enviromental, over the time. These new approach are applied generally in medicine, engineering and others area, and are excellent tool to have better explanation about sigatoka behaviour.

The incubation period determined experimentally in this study was similar to values found by other authors, such as 13 to 31 days (Marín et al., 2003) and 24.8 days at 30 ° C (Rocha, 2012). The occurrence of the disease was not influenced by the acclimation, growth system nor leaf age, confirming that the disease occurs in both shaded and non-shaded plants. However, the data show differences in the time for appearance of lesions in the different growth systems.

The longer incubation period for shaded grown plants indicates that illuminance interferes in fungal penetration and colonization. In fact, laboratory experiments (Santana-Filho et al., 2012) showed that illuminance decreases mycelial growth and spore germination. The explanation for the longer incubation is the possible lower production of phytotoxins and other pathogenesis-related compounds light-dependent, as suggested by Stierle et al. (1991), Daub et al. (2005) and Beltrán-García et al. (2014). On shade these compounds are not produced, affecting the epidemiological parameters penetration and colonization. Nevertheless, one study showed that there is no relation between light and phytotoxins produced by *Pseudocercospora fijiensis* (Cruz-Cruz et al., 2009). Further studies need to be conducted to shed light on this important aspect of the plant-pathogen relationship.

The effect of stomata density and infection by *P. musae* is not well understood. In this study, the leaves with high stomata density had short incubation time as the results showed by Craenem et al. (1997) that found negative correlation between Black Sigatoka incubation time and stomata density on polyploid hybrids.

The time for the appearance of the first sporulated lesions (latency) was similar to data found by other authors. For example, 46 days at 20°C, 32 days at 24°C and 33 days at 30°C in plantations of Saquarema variety, Cavendish (AAA) subgroup located in Minas Gerais, Brazil (Rocha, 2012), and 26 to 42.5 days for varieties Nanicão e Prata Anã in plantations of Bahia, Brazil (Cordeiro, 1997). Santana-Filho et al. (2012) show that low amounts of light reduce sporulation of *P. musae*, corroborating data from Sepúlveda et al. (2009).

The time to reach severity indices 4 and 6 was decreased in non-shaded grown plants, which is in agreement with reports affirming that growing banana in the shade may reduce the damage caused by black sigatoka (Norgrove, 1998, Norgrove et al. 2012, Norgrove & Hauser, 2012).

The infectious period is shorter in plants grown in the sun, but these plants produce more lesions and more spores as a result of the illuminance effect. In summary, illuminance influences all stages of the pathogen’s life cycle, and this study shown that from acclimatization, age from leaves and growth system, the last one was the most important variable influencing disease development. Whitin one year, according to the data analyzed, we have 10,72 cycles of latency in the Sun and 7,63 cycles in the shade condition. The same situation was found to severity 4 (5,49 cycles/year in the sun; 4,60 cycles/year in the shade) and severity >6 (1,99 cycles/year in the sun; 1,65 cycles/year in the shade). The data analyzed shows complete infection periods to 89% of plats in the sun in 60 days, so then in a year we could have 6 complete cycles for 89% of plants. In the shade conditions only 6% of plants could be have the same situation described above.

Differently from other studies, we reported the risks for the occurrence of different epidemiological events and the exact time they happened. The data may be used in the construction of parameters to model the temporal dynamics of yellow Sigatoka disease, which remains the most important foliar disease in Brazil. Our studies have shown that crop systems with shaded enviromental as agroforestry systems is better to banana cultivation because is the best cultural way to control the pathogen reproduction and dissemination.

## ACKNOWLEDGEMENTS

The authors are grateful to CNPq for their research grant. We also thank Mr Francisco Paulo dos Santos Souza (Embrapa Cassava&Fruits) for his technical assistance. DMSF is indebted to CAPES for scholarship in the master degree and Master Program in Agricultural microbiologist of UFRB for study opportunity. FFL is indebted to CNPq for his research grant.

